# Transcutaneous penetration of a single-chain variable fragment (scFv) compared to a full-size antibody: potential tool for Atopic Dermatitis (AD) treatment

**DOI:** 10.1101/2021.01.29.428747

**Authors:** Audrey Baylet, Raoul Vyumvuhore, Marine Laclaverie, Laëtitia Marchand, Carine Mainzer, Sylvie Bordes, Brigitte Closs-Gonthier, Laurent Delpy

## Abstract

**Background:** Currently, several biologics are used for the treatment of cutaneous pathologies such as atopic dermatitis (AD), psoriasis (PSO) or skin cancers. The main administration routes are subcutaneous and intravenous injections. However, little is known about antibody penetration through the skin.

**Objectives:** The aim was to study the transcutaneous penetration of a reduced-size antibody as a single-chain variable fragment (scFv) compared to a whole antibody (Ab) and to determine its capacity to neutralize an inflammatory cytokine involved in AD such as human interleukin-4 (hIL-4).

**Methods:** Transcutaneous penetration was evaluated by *ex vivo* studies on tape-stripped pig ear skin. Antibody visualization through the skin was measured by Raman microspectroscopy. In addition, hIL-4 neutralization was studied using two 2D models. First, embryonic alkaline phosphatase (SEAP) secretion by HEK-Blue™ IL-4/IL-13 cells, proportional to hIL-4 cells stimulation, was quantified by OD 620 nm measurement in presence or absence of an anti-hIL4 scFv or Ab. Then, normal human keratinocytes (NHKs) were stimulated with polyinosinic-polycytidylic acid (poly I:C) +/− hIL-4 and treated with anti-hIL4 scFv. Human Interleukin-8 (hIL-8) concentrations were determined in culture supernatants by ELISA.

**Results:** After 24h of application, analysis by Raman microspectroscopy showed that scFv penetrated into the upper dermis while Ab remained on the *stratum corneum*. In addition, the anti-hIL4 scFv showed better efficiency compared to Ab, with a neutralization percentage at 200 nM of 68% and 47%, respectively, in the HEK-Blue™ IL-4/IL-13 model. hIL-8 dosage in stimulated NHKs supernatants revealed that addition of scFv induced a dose-dependent hIL-4 neutralization.

**Conclusions:** scFv penetrates through to the upper papillary dermis while Ab remains on the surface. The anti-hIL4 scFv neutralizes its target effectively in two 2D models suggesting its potential use as topical therapy for AD.

## Introduction

Currently, immunotherapy is used for the treatment of several skin pathologies such as atopic dermatitis (AD), psoriasis (PSO) and cancers, mainly via subcutaneous or intravenous routes. Finding new antibody delivery methods is a challenge and different technologies have emerged in recent years such as microneedle patches, nanoparticles, liposomes or gel formulations (1). In addition, several modified antibodies are under development to treat cutaneous diseases, but at present, only certolizumab pegol, a PEGylated anti-tumor necrosis factor α antigen-binding fragment (Fab) has been approved (2013) for the treatment of PSO (1). AD is an inflammatory skin disease with a high and constantly increasing prevalence worldwide (1-3% of adults and 15-20% of children) (2). This pathology is characterized by a skin barrier default and a Th2 phenotype with secretion of several proinflammatory cytokines such as interleukin-4 (IL-4), IL-5 and IL-13. Among them, IL-4 is particularly involved in immunoglobulin (Ig) class switching to IgE (3). Different cell populations express the high affinity IgE receptor (FcεRI) and immune complex phagocytosis by these cells leads to T lymphocytes activation (4). Until now, only one immunotherapy treatment targeting the IL-4/IL-13 common receptor, IL-4 receptor alpha (IL-4Rα), has been approved for AD. Thus, developing a new topical immunotherapy treatment is a real challenge that would improve patients quality of life.

At present, there is no evidence in the literature proving that a reduced-size antibody could penetrate the skin more easily than a whole antibody. Thus, the first aim was to compare the passage of a single chain variable fragment (scFv) (~25 kDa) through a damaged skin, mimicking the poor barrier function of atopic skin, compared to a full-size monoclonal antibody (Ab) (~150 kDa). In addition, we compared if a scFv could neutralize human IL-4 (hIL-4) more effectively than an Ab in two 2D culture models.

## Methods

Briefly, *ex vivo* experiments were performed on pig ear skin. After shaving, ears were tape-stripped 25 times (Clinical&Derm, U.S.A.) to obtain the altered skin condition. Then, 8 mm diameter punches were treated with 5 µl phosphate buffered saline (PBS, Corning, U.S.A.), and 11 µM anti-CD44 scFv (Creative Biolabs, U.S.A.) or anti-CD44 Ab (Thermofisher Scientific, U.S.A.). Skin samples were placed at 37°C for 6h or 24h and cryopreserved until Raman microspectroscopy analysis.

Three 14 μm thick sections were used per sample and three Raman acquisitions were read for each section totaling nine measures per sample. Briefly, Raman images were obtained using a confocal Raman microspectrometer (Horiba Jobin Yvon, France) operating with a 660 nm laser. Labspec 6 software (Horiba Jobin Yvon, France) was used for acquisition and data pre-processing. Raman maps were recorded in an area of X: 10-μm /Y: 150-μm with 5 μm step size in the XY directions. Raman spectra were acquired in the 400 to 2300 cm^−1^ spectral range. To prevent background noise, Raman spectra were smoothed using a 2^nd^ order Savitzky-Golay type filter, baseline corrected using a 7^th^ order polynomial function, and spectra with an intensity under 400 cts in the 1530-1730 cm^−1^ range were suppressed. Next, Raman spectra were cropped to keep only the 400-785 cm^−1^ range of interest, which was used for vector normalization. Finally, fitting (unmixing) by classical least squares with a non-negativity constraint (NCLS) was used to estimate the contribution of the various skin components leading to the images reflecting the distribution of the specific scFv or Ab through the skin cryo-sections.

HEK-Blue™ IL-4/IL-13 cells (InvivoGen, France) were cultured according to manufacturer’s instructions. Briefly, 50 400 cells were seeded in 96-well plates. Then, cells were stimulated with 10 ng/ml hIL-4 (Miltenyi Biotec, Germany) and treated with anti-hIL4 scFv (Creative Biolabs, U.S.A.) or Ab (Biotechne, U.S.A.) at different concentrations. After 24h, QUANTI-Blue™ substrate was added to supernatants. After 2h30 at 37°C, SEAP levels were measured at 620 nm using a spectrophotometer.

Normal human keratinocytes (NHKs) were isolated from human skin samples from healthy donors (*n* = 4) undergoing medical surgery. The skin was collected after written informed consent from the donors and institutional approval. NHKs were stimulated with polyinosinic-polycytidylic acid (poly I:C) (2 μg/ml, Sigma, U.S.A.) +/− hIL-4 (25 ng/ml, Peprotech, U.S.A.) and treated with anti-hIL4 scFv at different concentrations. After 24h, supernatants were harvested to evaluate hIL-8 secretion by enzyme-linked immunosorbent assay (ELISA) according to manufacturer’s instructions (R&D Systems, U.S.A.). Cell protein concentrations were determined using the BCA Protein Assay kit (Thermofisher Scientific, U.S.A.) according to manufacturer’s instructions. hIL-8 secretion was normalized to total protein amounts.

Statistical analysis was performed using Prism GraphPad software (GraphPad Software, U.S.A.). Significant differences between samples and control were evaluated by Student’s *t*-test. *P* values < 0.05 were considered significant.

Some detailed protocols are available in the Supporting Information.

## Results

Three distinct peaks from the skin signal, corresponding to the scFv and Ab specific Raman signatures, were identified in the spectral range between 400 and 785 cm^−1^ (Fig.1a). This first analysis was necessary to visualize antibody penetration through the skin. At 6h, both scFv and Ab were only observed in the *stratum corneum* (depth: 10 – 20 μm) (Fig. 1b). At 24h, scFv penetrated to a depth of 130 μm, corresponding to the upper papillary dermis while Ab remained on the surface (Fig. 1c). Results were obtained on altered skin samples. Hematoxylin and eosin staining (H&E) of pig ear skin showed an altered skin barrier function on tape-stripped samples due to removal of part of the *stratum corneum* (Supplementary Figure S1).

**Figure 1.**
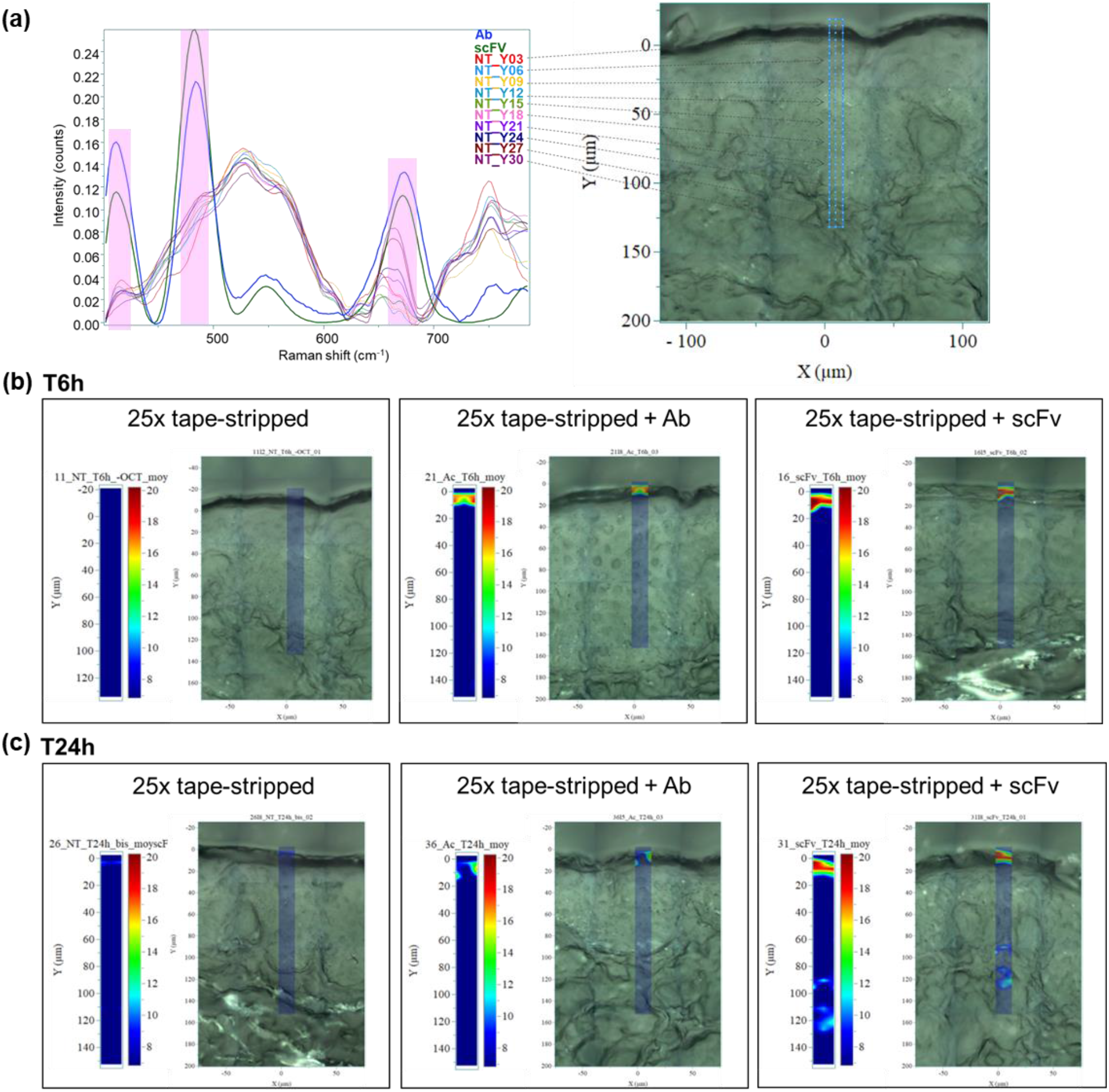
Visualization of transcutaneous antibody penetration by Raman microspectroscopy. **(a)** Single-chain variable fragment (scFv) and antibody (Ab) Raman signals (highlighted in pink) and skin Raman signals at different depths between 400 and 785 cm^−1^. **(b)** Quantification of scFv and Ab in damaged skin after 6h or **(c)** 24h of treatment.

Both the anti-hIL4 scFv and Ab showed significant dose-dependent neutralization of hIL-4 in the HEK Blue 2D model (Fig. 2a). In addition, at the maximal concentration of 200 nM, scFv neutralized an average of 68% of hIL-4 compared to 47% for Ab (Supplementary Table S1).

**Figure 2.**
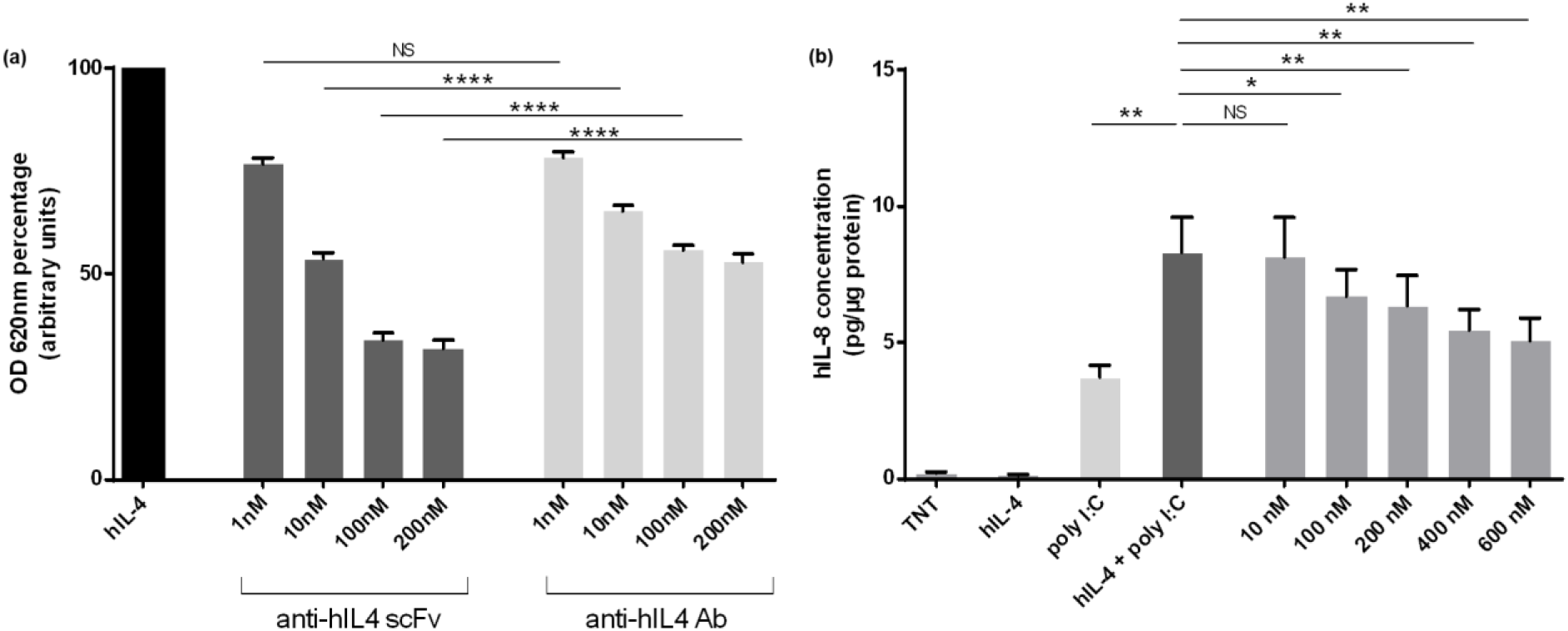
Human interleukin 4 (hIL-4) neutralization by single-chain variable fragment (scFv) on 2D models. **(a)** hIL-4 neutralization by scFv and Ab on HEK-Blue™ IL-4/IL-13 cells. Optical densities at 620 nm (OD 620nm) were determined and results were expressed in arbitrary units after normalization to the maximum value (100%) measured in cells stimulated with 10 ng/ml hIL-4. In each experiment and for each condition, cells were seeded in triplicates (*n* = 4 for 1, 10 and 10 nM doses; *n* = 3 for 200 nM dose). Results are expressed as mean +/− SEM. **(b)**Human interleukin 8 (hIL-8) quantification in supernatants of normal human keratinocytes (NHKs) stimulated with hIL-4 +/− polyinosinic-polycytidylic acid (poly I:C) (25 ng/ml and 2 μg/ml, respectively) and treated with anti-hIL4 scFv at different concentrations for 24h (*n* = 4). For the four donors, experiments were performed in duplicates. Results are normalized in relation to the quantity of total proteins per well and expressed as mean +/− SEM. (NS: non-significant; \**P* < 0.05; \*\**P* < 0.01; \*\*\**P* < 0.001; \*\*\*\**P* < 0.0001).

hIL-8 dosage of supernatants from NHKs stimulated with hIL-4 +/− poly I:C and treated with anti-hIL4 scFv, revealed a scFv dose-dependent decrease in four different donors (Fig. 2b). At 100, 200, 400 and 600 nM doses, a significant decrease in hIL-8 secretion was observed. ScFv neutralization ability against hIL-4 was calculated based on poly I:C induction of hIL-8. The hIL-4 + poly I:C was considered as the 0% of hIL-4 neutralization value, whereas poly I:C alone was considered as the 100% value (Table 1). At 600 nM scFv, the average percentage of hIL-4 neutralization reached 80%. In addition, anti-hIL4 Ab showed low neutralization efficiency in this 2D model (data not shown).

**Table 1.** hIL-4 neutralization by scFv in normal human keratinocytes (expressed as a relative percentage to poly I:C) Human interleukin 4 (hIL-4) neutralization by single-chain variable fragment (scFv) determined in normal human keratinocytes (NHKs) (*n* = 4) (expressed as a relative percentage to polyinosinic-polycytidylic acid (poly I:C)).

| Donors | poly I :C | hIL-4 + poly I :C | scFv 10 nM | scFv 100 nM | scFv 200 nM | scFv 400 nM | scFv 600 nM |
| --- | --- | --- | --- | --- | --- | --- | --- |
| D1 | 100 | 0 | -39 | 4 | 26 | 44 | 84 |
| D2 | 100 | 0 | 69 | 84 | 118 | 113 | 127 |
| D3 | 100 | 0 | 3 | 21 | 31 | 46 | 43 |
| D4 | 100 | 0 | -7 | 33 | 28 | 58 | 64 |
| Average (%) | 100 | 0 | 7 | 36 | 51 | 65 | 80 |
| Standard deviation (%) | 0 | 0 | 45 | 35 | 45 | 32 | 36 |

hIL-4: human interleukin 4. poly I :C : polyinosinic-polycytidylic acid. scFv: single-chain variable fragment. nM: nanomolar.

## Discussion

Until now, few studies assessing antibody penetration in the skin have been carried out. Among them, topical application of infliximab, an anti-TNFα Ab, has been described in patients with ulcers (5), and Flightless I (Flii) neutralizing antibodies (FnAb) have been applied in a murine model of epidermolysis bullosa acquisita (6,7). No comparative study examining the penetration of reduced-size versus whole antibodies has yet been performed, especially using Raman microspectroscopy, which is a relevant tool for antibody characterization. Indeed, this technique can be used to monitor post-translational modifications, degradation or aggregation (8–10). For several years, it has also been used to study the skin (11,12). To our knowledge, we are the first to track antibody passage through the skin using Raman microspectroscopy. Here, we showed in an *ex vivo* model of damaged skin that reduced size facilitated antibody penetration into the upper papillary dermis.

Only one antibody fragment has been approved for the treatment of skin pathologies. Recently, M1095, an anti-IL17A/F nanobody completed a phase 1 clinical trial for the treatment of moderate to severe PSO (13). Thus, antibody fragments could be potential tools for the development of new topical treatments for cutaneous diseases. Indeed, in recent years, innovative immunotherapy methods have emerged including microneedles, nanoparticles or liposomes (14–17). To date, only one immunotherapy treatment has been approved for AD. Several treatments targeting proinflammatory molecules including cytokines and their receptors are under development for AD management (18). First, we chose to target hIL-4, which is a key cytokine involved in AD physiopathology. In fact, it leads to B-lymphocytes class switching to IgE and naïve T cells differentiation into Th2 (19). Therefore, we decided to focus on the effect of hIL-4 neutralization on inflammatory human keratinocytes. We found a scFv dose-dependent decrease in hIL-8 secretion suggesting a key role for hIL-4 in inflammation. In addition, our results demonstrated the neutralization efficiency of anti-hIL4 scFv on HEK-Blue™ IL-4/IL-13 cells, which performed better than the whole antibody itself.

A limitation of our *ex vivo* model could be the lack of lesional AD human skin biopsies. Tape strips represent a simple and easy way to mimic barrier disruption, feature observed in the pathology. Nevertheless, this alteration does not reflect all structural changes of atopic skin, suggesting a different behavior of scFv penetration on patient’s skin that would need to be assessed. In the future, it would be interesting to evaluate the benefits of a topical anti-mouse IL-4 treatment on an *in vivo* AD model such as NC/Nga mice (20).

To sum up, we showed that reduced-size antibodies depict better penetration abilities than full-size antibodies and remain as effective to neutralize their targets. Reduced-size antibodies could be therefore potential relevant topical treatments for inflammatory skin diseases as AD.

## Acknowledgements

We would like to thank all the members of the UMR CNRS 7276 – INSERM U 1262 research unit for helpful discussions and comments and Jeanne Cook-Moreau for proofreading of the manuscript. We would also like to thank all the members of Silab R&D and more specifically Nathalie Solingeas and Elodie Brugère for their technical expertise. This work was supported by grants from Silab R&D. A.B. is a recipient of a Ph.D. fellowship Cifre from the French Ministry of Research.

The authors declare no competing financial interests.

---

**Figure S1.**
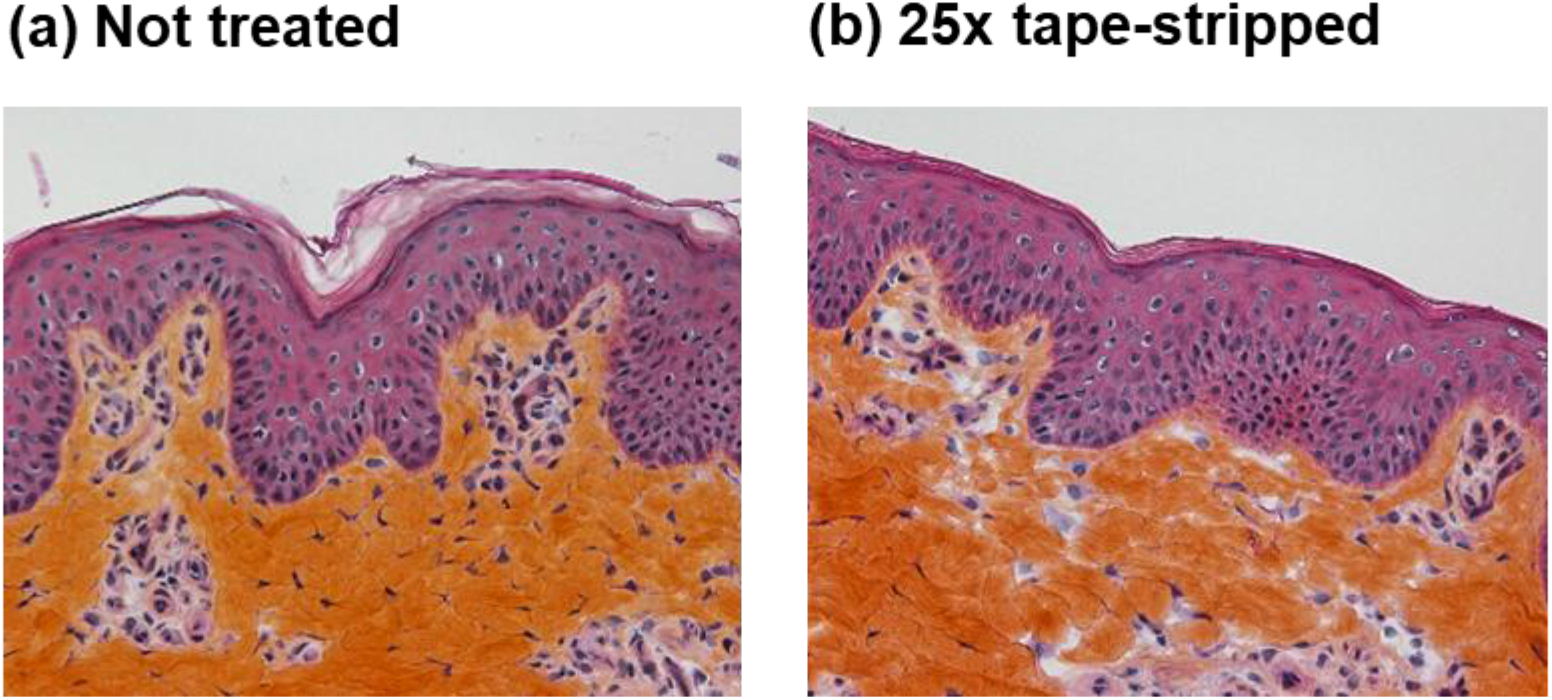
Representative images for hematoxylin and eosin staining (H&E) of pig ear skin samples (a) with no treatment or (b) 25 times tape-stripped to mimic damaged skin.

**Table S1:** hIL-4 neutralization by scFv in HEK-Blue™ IL-4/IL-13 cells (expressed as a relative percentage to OD620nm) Human interleukin 4 (hIL-4) neutralization by single-chain variable fragment (scFv) determined on HEK-Blue™ IL-4/IL-13 cells (expressed as a relative percentage to optical density at 620 nm (OD620nm)).

|  | scFv (nM) |  |  |  | Mab (nM) |  |  |  |  |
| --- | --- | --- | --- | --- | --- | --- | --- | --- | --- |
|  | 1 | 10 | 100 | 200 | 1 | 10 | 100 | 200 | No treatment |
| Exp. 1 | 27 | 49 | 69 |  | 24 | 31 | 41 |  |  |
| Exp. 2 | 24 | 49 | 72 | 75 | 23 | 41 | 51 | 53 | 0 |
| Exp. 3 | 18 | 41 | 61 | 64 | 24 | 32 | 44 | 44 | 0 |
| Exp. 4 | 24 | 46 | 62 | 65 | 16 | 34 | 42 | 45 | 0 |
| Average (%) | 23 | 46 | 66 | 68 | 22 | 35 | 44 | 47 | 0 |
| Standard deviation (%) | 4 | 4 | 5 | 6 | 4 | 4 | 4 | 5 | 0 |

hIL-4: human interleukin 4. IL: interleukin. scFv: single chain variable fragment. Exp.: experiment. nM: nanomolar.

## Supporting information

### Supplementary material

#### HEK-Blue™ IL-4/IL-13 cells

IL-4/IL-13 binding to their common receptor subunit, IL-4 receptor alpha (IL-4Rα), leads to signal transducer and activator of transcription 6 (STAT6) phosphorylation by Janus kinases (JAK) and its translocation in the nucleus to activate target genes. HEK-Blue™ IL-4/IL-13 cells (InvivoGen, France) specifically express the reporter gene secreted embryonic alkaline phosphatase (SEAP) under the control of a specific promotor fused to several STAT6 binding sites. After stimulation with hIL-4, SEAP was detected in culture supernatants using QUANTI-Blue™ substrate. In each experiment and for each condition, cells were seeded in triplicates (*n* = 4 for 1, 10 and 10 nM doses; *n* = 3 for 200 nM dose).

#### Normal human keratinocytes (NHKs)

NHKs were cultured in Keratinocyte serum-free medium (KSFM) (Thermofisher Scientific, U.S.A.) supplemented with human recombinant epidermal growth factor (rEGF) and bovine pituitary extract (BPE) at the time of use and were harvested at third-passage by trypsin digestion. At day 0, 40 000 cells per well were seeded in 24-well plate in KSFM rEGF BPE. At 80% confluence, NHKs were stimulated with polyinosinic-polycytidylic acid (poly I:C) (2 μg/ml, Sigma, U.S.A.) +/− hIL-4 (25 ng/ml, Peprotech, U.S.A.) and treated with anti-hIL4 scFv (10, 100, 200, 400 and 600 nM) diluted in 500 μl KSFM rEGF BPE. For each condition, experiments were carried out in duplicates. After 24h, supernatants were harvested to evaluate hIL-8 secretion by enzyme-linked immunosorbent assay (ELISA) according to manufacturer’s instructions (R&D Systems, U.S.A.). Cells were lysed by adding 300 μl of NaOH 0.1N and cellular proteins concentration was determined using the BCA Protein Assay kit (Thermofisher Scientific, U.S.A.) according to manufacturer’s instructions.

